# Selective inhibition of fibroblast-specific Domain Discoidin Receptor 1 (DDR1) reduces collagen deposition and modulates fibroblast-specific cytokine release within the breast microenvironment

**DOI:** 10.1101/2023.08.11.552937

**Authors:** Daniel E.C. Fein, Nicole Traugh, Gat Rauner, Youssof Mal, Colin Trepicchio, Charlotte Kuperwasser

## Abstract

Fibroblasts are a major cell type within breast microenvironment which play key roles in tissue remodeling during the processes of normal development, injury, and malignancy. During wound healing and tumorigenesis, fibroblasts facilitate production and degradation of the extracellular matrix and produce inflammatory mediators which act as immune regulators. Domain Discoidin Receptor 1 (DDR1) is a cell surface tyrosine kinase receptor expressed by epithelial and stromal cells which is activated by collagen. In the breast, DDR1 expression and activity has been implicated in the development of fibrosis as well as chemotherapy resistance. We set out to examine whether selective inhibition of DDR1 would modulate fibroblast immunomodulatory function to generate an immune-permissive breast microenvironment and reduce stromal desmoplasia. In vivo, DDR1 inhibition resulted in mammary fibroblast tissue remodeling, reduced collagen deposition, and changes in immunomodulatory cytokine expression. Furthermore, DDR1 inhibition was associated with increased CD45.2+ immune cell infiltration and reduced Ly6G+/Ly6C− neutrophil infiltration. Mechanistically, we developed an ex-vivo 3D collagen hydrogel model of desmoplasia to study the effects of DDR1 inhibition on the expression of immune modulating factors and fibroblast functions and features. We found that DDR1 regulates the expression and secretion of key immunomodulatory cytokines (IL-6, IL-8, and MCP-1). Collectively these findings suggest that breast fibroblast-specific DDR1 mediates collagen deposition and immunomodulatory function within the mammary gland and warrants further investigation as a potential target for fibroblast-modulating therapy in benign and neoplastic breast disorders.

## Introduction

The fibroblast is the principal cell that synthesizes collagen and extracellular matrix of the mammary stroma. Now considered a major participant in the mammary immune response, fibroblasts have emerged as a new target for treatment of benign and neoplastic breast disorders given their contribution to desmoplasia, tumor growth, metastasis, and suppression of the anti-tumor immune response^1–4^. Fibroblasts activate in response to stress and pathological conditions by transitioning to a contractile phenotype as such^1,5^. This process is referred to as desmoplasia.

In the breast, cancer-associated fibroblasts (CAFs) are predominantly formed from conversion of tissue-resident mammary fibroblasts through varied mechanisms including increasing extracellular matrix (ECM) stiffness, ECM composition, hypoxia, and overproduction of activating signals such as TGFβ and IL-6^1,6–8^. The clinical experience in targeting CAFs has been limited and in some instances has resulted in paradoxically increased tumor aggressiveness^2,9,10^. As a result, there has been a call for further understanding of the relation of fibroblast markers to function as well as to prioritize strategies aimed to reprogram CAFs rather than ablate them^11^.

Domain Discoidin Receptor 1 (DDR1) is a cell surface tyrosine kinase receptor expressed by epithelial and stromal cells which is activated by collagen within the breast microenvironment^12–14^. DDR1 has been identified as a key mediator of the stromal-epithelial interactions during ductal morphogenesis in the human mammary gland^15,16^. In breast cancer, DDR1 expression has been associated with desmoplasia, chemotherapy resistance, tumor invasiveness, and metastasis of yet unclear mechanisms^13,17–20^. Most recently, tumor DDR1 has been implicated in immune cell exclusion through a mechanism of collagen fiber realignment within the TME^21^. The effects of pharmacologic targeting of DDR1-dependent stromal-epithelial interactions on the desmoplastic response and immune cell composition remain unclear. In this study, we set out to study how desmoplasia may affect an immunological response by examining whether selective inhibition of DDR1 would affect fibroblast immunomodulatory function. In doing so, we found that inhibiting collagen signaling can generate an immune-permissive breast microenvironment and reduce stromal desmoplasia within the benign mammary gland.

## Results

### DDR1 inhibition reduces collagen deposition and modulates expression of inflammatory cytokines and markers of ECM remodeling

Fibroblast release of immunomodulatory cytokines has been implicated as a driver of the desmoplastic response through numerous cell-dependent and independent interactions^1^. These interactions may be context dependent, in that the stromal secretome may be influenced by the composition and stiffness of the surrounding ECM^1^. To study the role of mammary fibroblasts connection between role of DDR1 on the mammary stroma, extracellular matrix remodeling, cytokine profile, and immune cell composition, we treated 12-week-old C57Bl/6 female mice with an inhibitor of DDR1 (Fig 1A). Mice were administered an orally bioavailable selective DDR1 inhibitor (DDRi) once daily for 14 days, after which thoracic and inguinal mammary glands (#2-3 and #4-5, respectively) were collected and examined. Picrosirius red staining of inguinal mammary glands after treatment showed a reduction in collagen content within the mammary tissue (Fig 1B, 1C). Gene expression analysis showed reduced expression of markers of ECM remodeling in the DDR1i group including collagen type 1 (*Col1a1*), collagen type 4 (*Col4a1)*, and *Mmp9* (Fig 1E).

**Figure 1 -.**
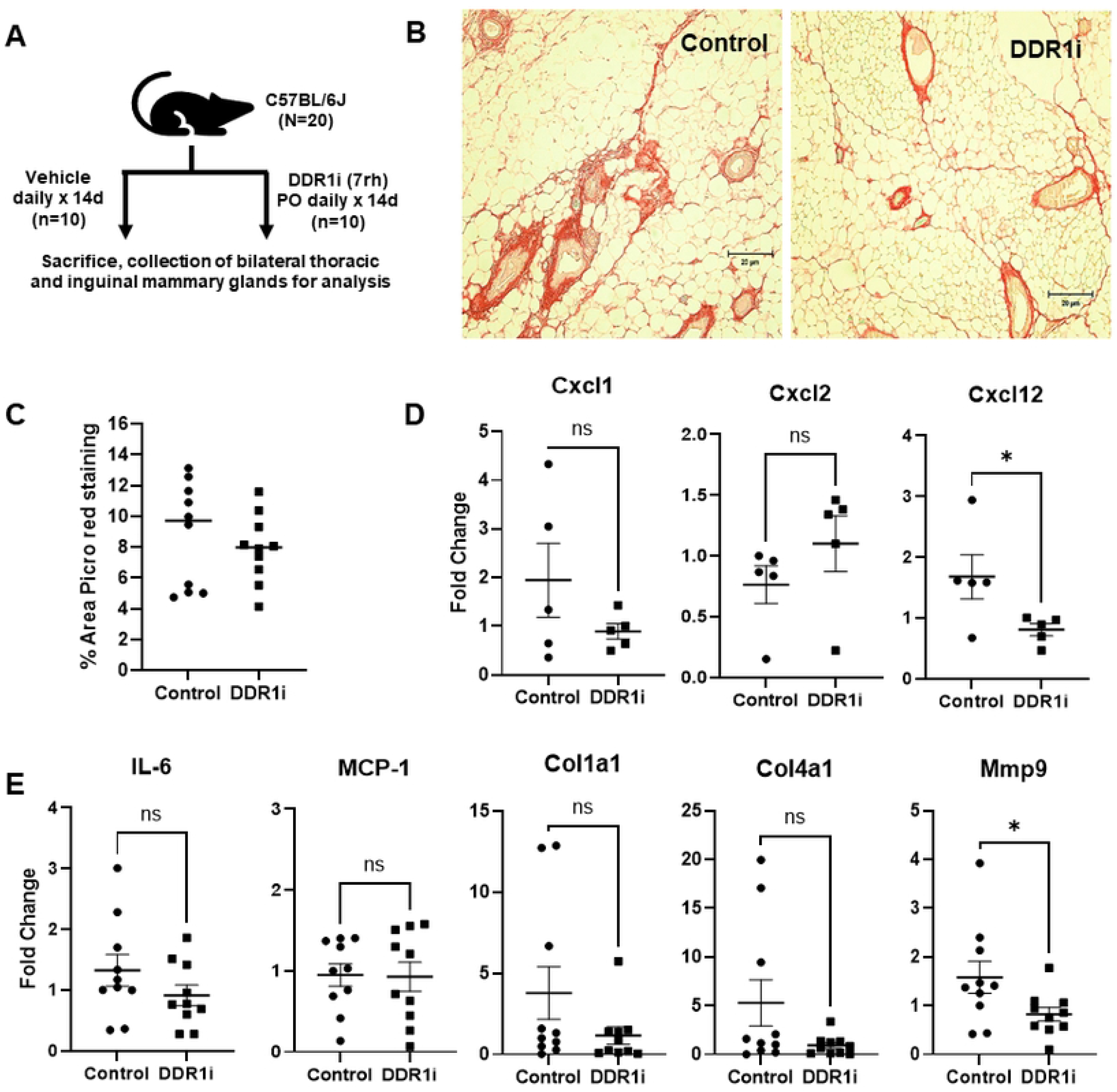
Effect of selective DDR1 inhibition (DDR1i) *in vivo* on collagen deposition, extracellular matrix (ECM) remodeling, and inflammatory cytokine expression within the mammary gland. Twenty 12-week-old C578L/6J female mice were randomized in 1:1 fashion to receive either DDR1i (7rh 25 mg/kg once daily) or vehicle (1% carboxymethylcellulose, 0.25% Tween 80) via oral gavage for 14 days, followed by sacrifice and immediate collection of mammary glands (A). Picrosirius red staining was performed on paraffin-embedded inguinal mammary glands (B), followed by quantification of staining to assess collagen deposition (C). Total RNA was extracted from thoracic mammary glands followed by RT­ qPCR gene expression analysis to assess the relative expression of relevant inflammatory chemokines, cytokines, and markers of ECM remodeling (D,E,). * p <0.05, ns - not statistically significant.

Given the reduction in immunomodulatory cytokine expression and collagen content in mammary tissues, we examined the immune cell composition of the mammary gland. Using an established immunophenotyping panel^28^, multiplexed flow cytometry of inguinal mammary glands was performed to assess the innate and adaptive immune cell compositions (Fig S1, Fig S2). We observed trends in increased CD45.2+ hematopoietic cell percentage in the DDR1i group (Fig 2A). Also observed in the DDR1i group were trends in reduced CD11b+ myeloid and CD11b+Ly6G+Ly6C− neutrophil populations (Fig 2A). The relative percentage of CD11b+Ly6G-Ly6C− macrophages were increased. The total percentage of CD19+TCRβ-B cells and CD19-TCRβ+ T cells were increased in the DDR1i group (Fig 2B). Consistent with findings in the mammary stroma, the spleen also showed a significant reduction in the CD11b+Ly6G+Ly6C− neutrophil population and a significant increase in CD11b+Ly6G-Ly6C− macrophage population (Fig 3A). Associated with reduction in neutrophil population, we observed reduced gene expression of potent chemokines *Cxcl1* and *Cxcl12* (Fig 1D) in the mammary gland. In association with the trends of increased macrophage infiltration seen (Fig 2A, 3A), trends in increased *Cxcl2* expression were also observed in the mammary glands of the DDR1i group (Fig 1D).

**Figure 2 -.**
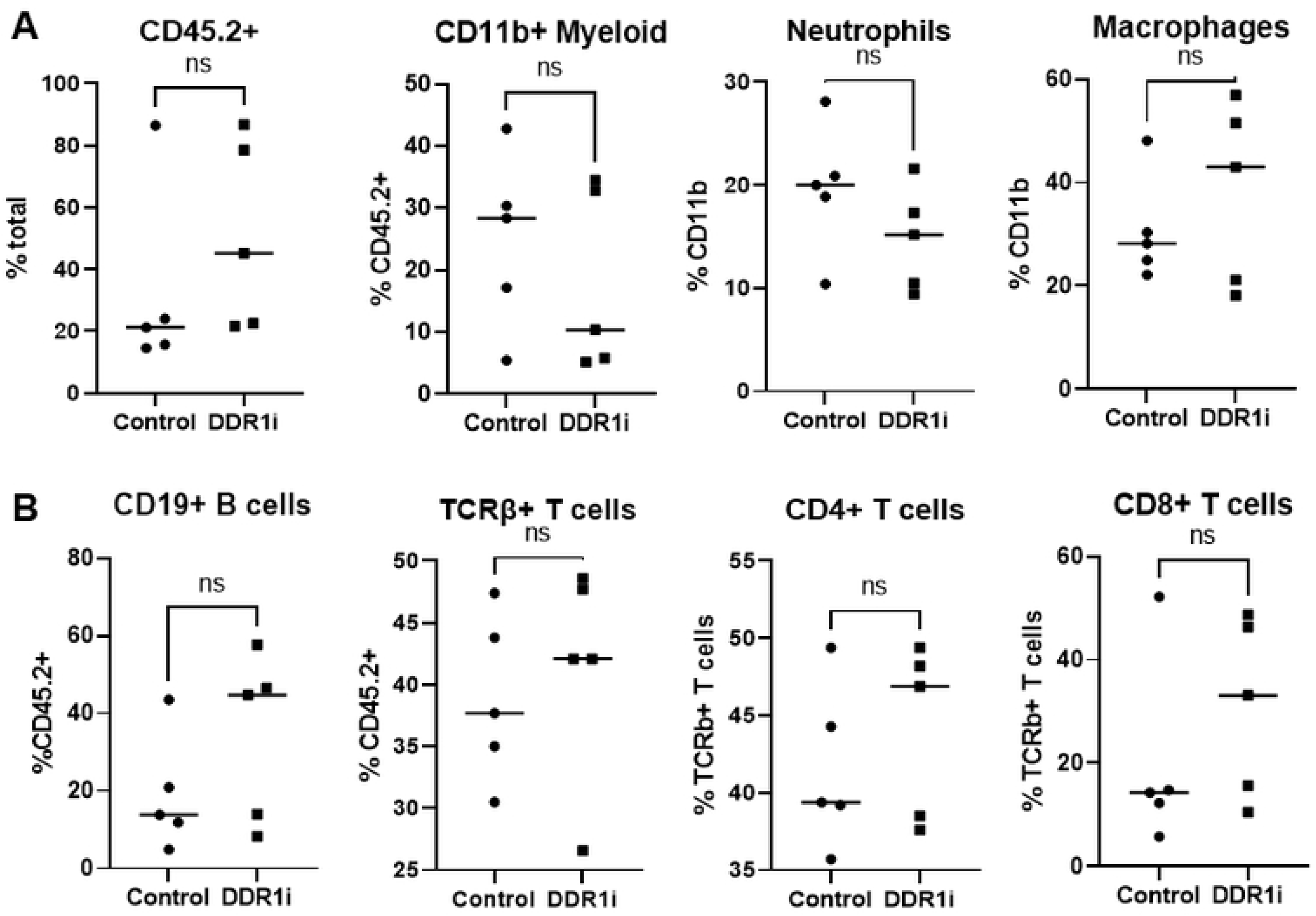
Selective DDR1i *in vivo* modulates the immune cell composition of the mammary gland. Murine inguinal mammary glands were harvested immediately after sacrifice and used to form single cell suspensions through a 1-hour collagenase/hyaluronidase/TrypLE digestion method. This was followed by immediate assessment of myeloid (A) and lymphoid (B) cell populations using a multiplexed flow cy1ometry approach (Figure S1, S2). ns - not statistically significant.

**Figure 3 -.**
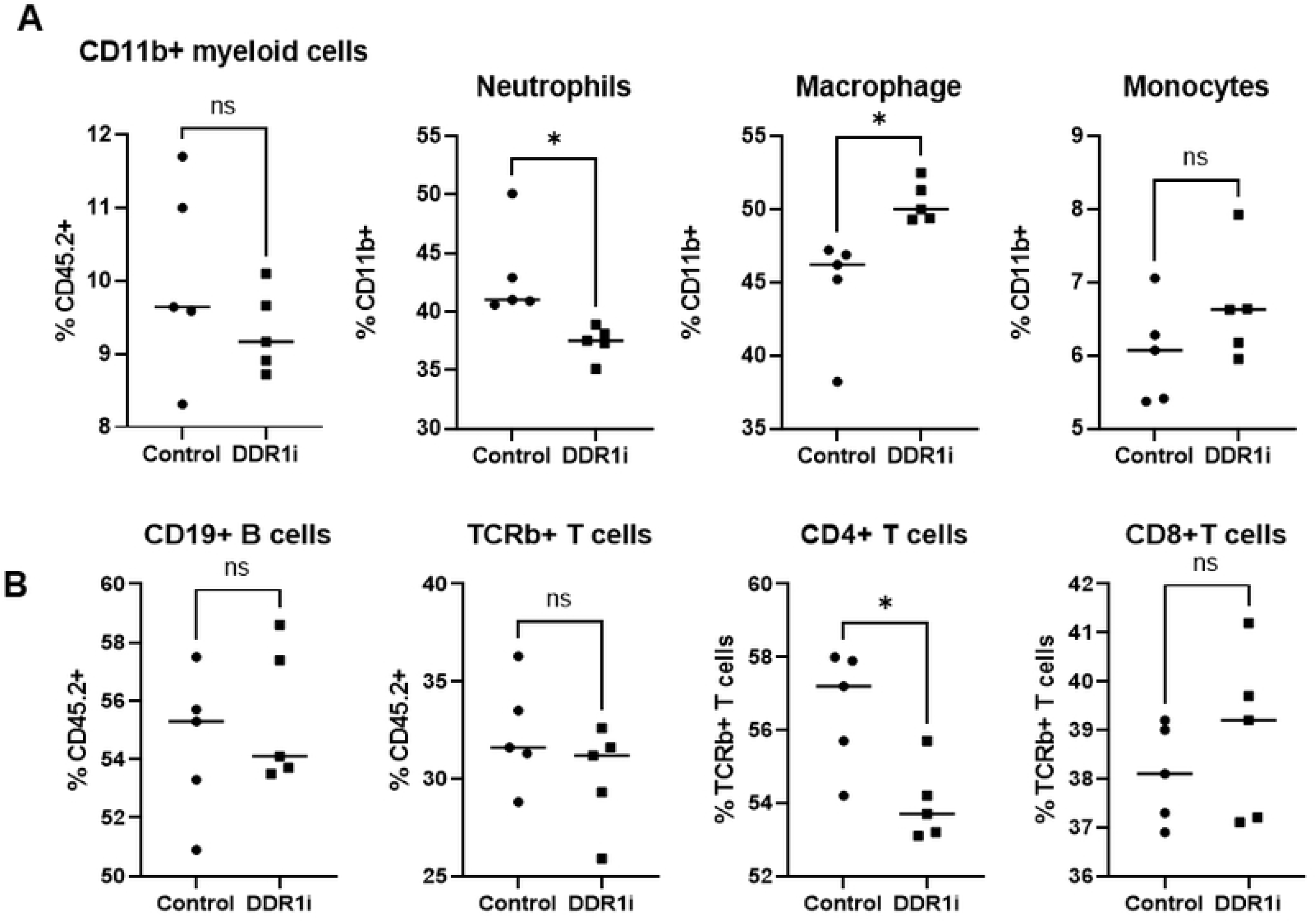
Selective DDR1i *in vivo* modulates the immune cell composition of the spleen. Murine spleens were harvested immediately after sacrifice and used to form single cell suspensions through a mechanical digestion method. This was followed by immediate assessment of myeloid (A) and lymphoid (B) cell populations using a multiplexed flow cytometry approach (Figure S1, S2). * p <0.05, ns - not statistically significant.

### Collagen concentration and cell density have an important role in the contraction and expression of immune modulating factors by mammary fibroblasts

To study how DDR1i leads to changes in stromal collagen, we characterized the effect of ECM and cell density on the expression of key cytokines previously identified as drivers of tumor growth, metastasis, and modulators of the immune microenvironment^22–25^. Immortalized human breast tissue fibroblasts obtained from reduction mammoplasty tissue (RMF-EG fibroblasts^31^) were seeded into 3D collagen-I gels at increasing stiffness and cell density (Fig 4A). Fibroblasts in low stiffness gels exhibited increased contraction (Fig 4A). Additionally, fibroblasts in the fully contracted gels were observed to upregulate pro-inflammatory cytokines including *IL6* and *IL1b* (Fig 4B). By increasing cell density, a modest increase in collagen gel contraction was observed (Fig 4C). However, this modest increase in gel contraction was associated with marked upregulation of *IL6* and *IL1b* (Fig 4D). This may suggest that the pro-inflammatory secrotome of mammary fibroblasts is regulated significantly by both cell density as well as ECM stiffness and composition. Given these observations, we hypothesized that interruption of cell-ECM interactions through selective DDR1 inhibition could reprogram mammary fibroblast ECM remodeling and immune modulating functions.

**Figure 4 -.**
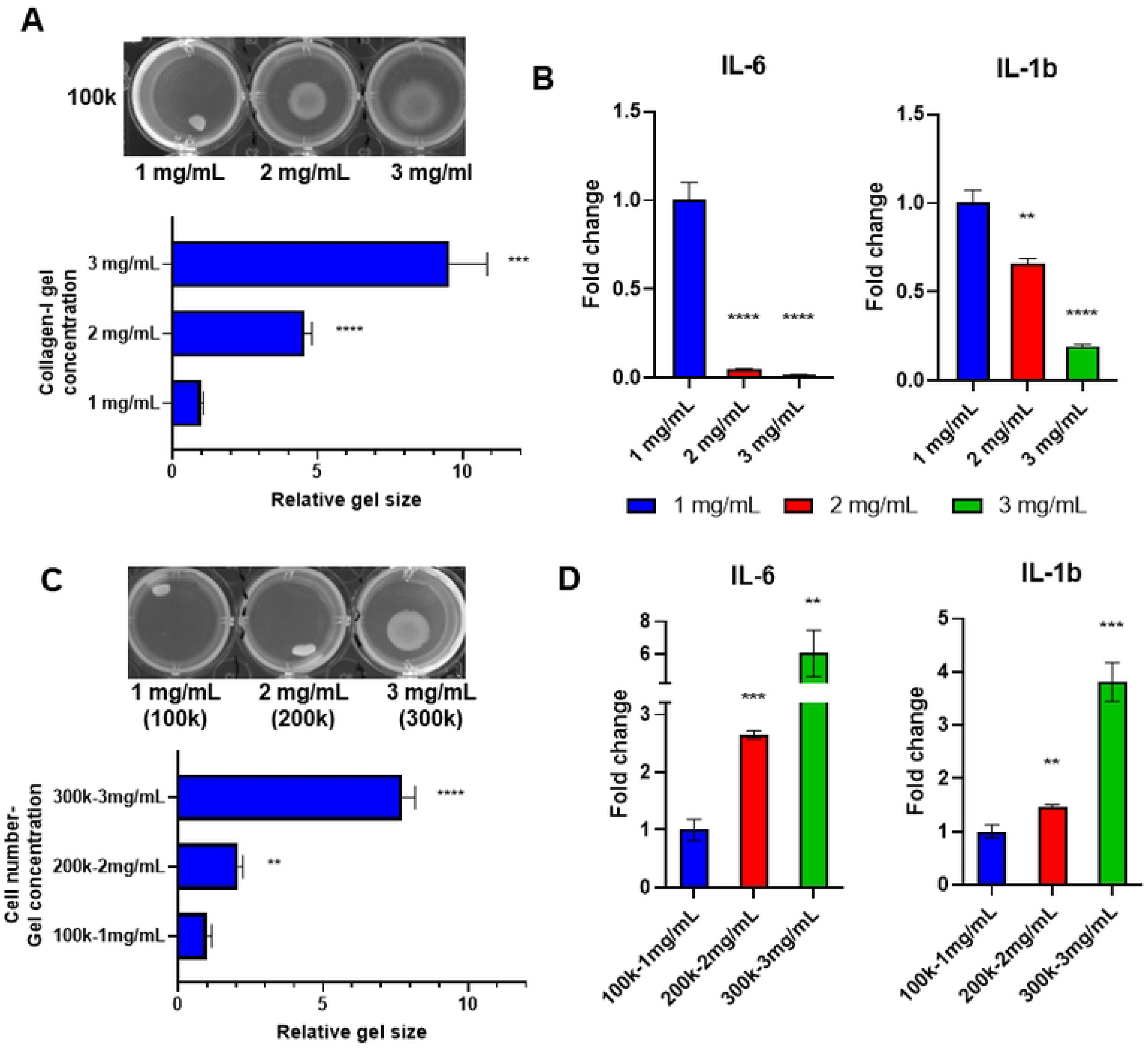
lmmunomodulatory cytokine expression of human mammary fibroblasts *in vitro* is influenced by collagen and cell density. Human mammary fibroblasts (RMF-EG) were seeded into 3D collagen hydrogels at increasing collagen concentration, with increased gel contraction observed at lower collagen gel concentration (A). RNA was isolated from fibroblast-laden gels followed by assessment of relative gene expression of pro-inflammatory cytokines via RT-qPCR analysis, with observation of increased *IL-6 and IL-1b* expression in gels with lowest collagen concentration/highest gel contraction (B). Increasing fibroblast cell concentration only partially improves gel contraction (C), while *IL-6 and IL-1b* expression is increased. * p<0.05, ** p<0.01, *** p<0.001, **** p<0.0001.

### Selective DDR1 inhibition modulates human fibroblast expression of cytokines and markers of ECM remodeling in vitro

We next sought to characterize the changes in fibroblast cytokine expression profile induced by DDR1i. RMF-EG were seeded into collagen gels with or without the addition of selective DDR1 tyrosine kinase inhibitors^26,27^. With DDR1i we observed significant reduction in RMF-EG-mediated collagen gel contraction (Fig 5A) in association with reduced DDR1 phosphorylation (Fig S3). Reduced expression of CAF markers *αSMA* and *FAP* were seen with DDR1i, in addition to marked reduction in *IL6* expression (Fig 5B). Conditioned media from RMF-EG fibroblasts cultured in 3D was collected for multiplex human cytokine array analysis. We identified elevated expression of cytokines IL-6, IL-8, and MCP-1 which were reduced with DDR1i (Fig 5C). No significant difference in IL-1b was observed (Fig 5C). Associated with reduced cytokine secretion, reduced mRNA expression of *IL6, IL8,* and *MCP1* with DDR1i was observed (Fig 5C,D). Interestingly, we observed that the relative reduction in mRNA expression of *IL6, IL8,* and *MCP1* was enhanced in RMF-EG grown in 3D collagen gels compared to those grown in 2D. In assessing genetic markers of ECM remodeling, we observed upregulation of collagen type 1 (*Col1a1)* and matrix metalloproteases (*Mmp9*) in RMF-EG cultured in the presence of collagen; DDR1i-induced downregulation of *Col1a1* and *Mmp9* was seen only in the 3D context (Fig 5D). Additionally, when stimulated to a myofibroblast phenotype with prolonged TGFβ1 treatment, RMF-EG fibroblasts were noted to have significant downregulation of *IL6* and *αSMA* when cultured in either 2D or 3D (Fig S5). To correlate these findings with our *in vivo* studies, we assessed mRNA expression of key cytokines in the mouse mammary gland, observing trends reduction of *IL6*, collagen, and matrix metalloprotease expression in the DDR1i treated mice compared with vehicle (Fig 1E).

**Figure 5 -.**
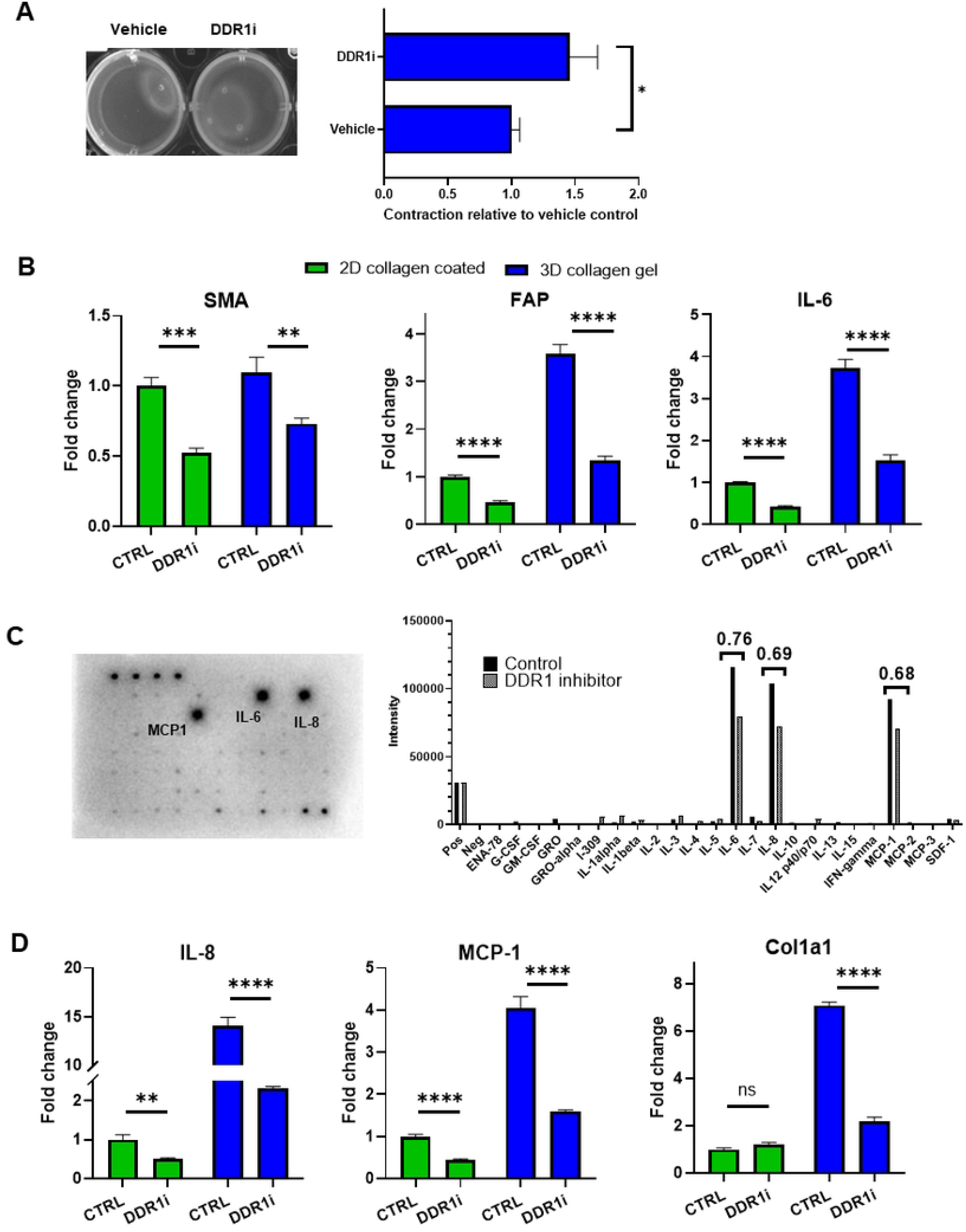
Selective DDR1i of human mammary fibroblasts cultured in 3D collagen hydrogels reduces expression of inflammatory cytokines and markers of fibroblast function. RMF-EG were cultured in 3D collagen gels in the presence of DDR1i (+) or vehicle (−) followed by assessment of collagen gel contraction [n=3 gels per condition] (A). Reduced gel contraction with DDR1i was observed (A). RMF-EG were cultured on collagen-coated tissue culture plate or in 3D hydrogel, followed by RNA isolation. RT-qPCR gene expression analysis demonstrated that DDR1 inhibition reduces expression of CAF markers *SMA, FAP* in the 2D or 3D context (B). *IL-6* was upregulated in RMF-EG cultured in 3D and downregulated with DDR1 inhibition (B). Cell supernatents from RMF-EG laden collagen gels were collected for multiplex human cy1okine array analysis. We identified high secretion of IL-6, MCP-1, and IL-8 which were reduced with DDR1 inhibition (C). To confirm this finding, RNA was isolated from RMF-EG cultured in 3D hydrogel followed by RT-qPCR gene expression analysis. RMF-EG in 3D hydrogels showed higher expression of *IL-8, MCP-1,* and *Col1a1* compared to 2D, subsequently reduced with DDR1 inhibition (D). * p<0.05, ** p<0.01, *** p<0.001, **** p<0.0001.

## Discussion

Investigations of fibroblast reprogramming in efforts to alter the disease course of fibrotic and malignant breast conditions have been limited by the 2D culture context. Using 3D culture models, one may more accurately emulate the composition and stiffness of the tissue microenvironment, factors known to significantly impact the proteome of supportive stromal cells^29^. Using a 3D collagen type I hydrogel model, we demonstrated the impact of breast fibroblast cell density, collagen density, and collagen gel contraction on the expression of potent immunomodulatory cytokines and subsequently the impact of interrupting stromal-ECM interactions through DDR1i. As stromal-collagen signaling through DDR1 has been implicated in the development of desmoplasia, treatment resistance, and tumor invasiveness, we hypothesized that modulation of the breast fibroblast secretome through DDR1 could reduce collagen deposition within the mammary gland.

In a DDR1 knockout (KO) animal model, it was observed that deletion of DDR1 reduced the growth of transplanted DDR1-intact murine mammary tumors and altered extracellular matrix remodeling of the tumor microenvironment^30^. Screening of the stromal fraction secretome identified IL-6 as a cytokine which was secreted in a DDR1-dependent manner, a mechanism subsequently implicated in accelerating tumor cell invasion and growth *in vivo*. These findings suggested that stromal cell-specific production of soluble factors such as IL-6 are functionally linked to DDR1 activity in the stromal compartment and are sufficient to accelerate breast cancer growth and aggressiveness^30^.

Accordingly, using small molecular DDR1 inhibitors, we observed downregulation and reduced secretion of breast fibroblast-specific IL-6 amongst other key immunomodulatory cytokines including IL-8 and MCP-1. We noted an enhanced effect of IL-6 reduction when fibroblasts were cultured in 3D collagen-I gels, suggesting a context-dependent mechanism of DDR1 signaling which is enhanced in a collagen-rich environment. Treatment of mice with DDR1i resulted in downregulation of IL-6 within the benign mammary gland. This finding was associated with downregulation of type 1 and 4 collagens as well as reduced collagen deposition. These findings support the existing evidence which have correlated DDR1 signaling activity with ECM-remodeling within the mammary gland.

A novel mechanism of tumor cell-mediated immune cell exclusion facilitated by DDR1-dependent collagen fiber realignment within the TME was recently uncovered^21^. In this study, DDR1-KO breast tumors implanted into DDR1-intact immunocompetent hosts were found to have increased CD4+ and CD8+ T cell infiltration with reduced tumor growth in association. This finding was not observed in immunodeficient hosts, suggesting a key role of DDR1 in regulating the adaptive immune system response to tumor growth. The binding of the DDR1 extracellular domain (DDR1-ECD) to collagen fibers was implicated in the enforcement of a stromal barrier contributing to the immune cell exclusion seen^21^.

We thus characterized the lymphoid and myeloid immune cell populations within the benign mammary glands of mice treated with or without DDR1i and observed increased infiltration of CD45.2+ immune cells, CD19+ B cells, and TCRβ+ T cells with DDR1i. These findings contribute to existing evidence supporting the role stromal-cell specific DDR1 signaling in regulation of the immune cell composition of the mammary gland. In addition to interruption of DDR1-ECD-dependent collagen realignment, our findings suggest that DDR1 inhibition may further modify ECM remodeling through reduced production of collagens and matrix metalloproteases.

This study had limitations. Our *in vitro* studies do not fully address the impact of DDRi on the modification of associated mechanotransduction signaling pathways (e.g. TNFα/NF-κB, integrin, Rho GTPase-activating protein signaling), although our preliminary studies have not suggested TNFα/NF-κB to be implicated (Fig S6). We did not assess the impact of DDR1i *in vivo* in the context of induced desmoplasia, however, the reductions in *αSMA* and *IL6* expression observed in the TGFβ-induced myofibroblast phenotype (Fig S5) suggest potent activity of DDR1i in that context. *In vivo*, we did identify similar alterations of immune cell populations within the spleen, however, we did not assess further the systemic effects of DDR1i as it relates to chemokine expression and collagen deposition within other tissues.

In summary, these findings suggest that disruption of fibroblast-ECM signaling through DDR1 inhibition may be a novel strategy to reprogram the desmoplasia-inducing and tumor-promoting features of mammary fibroblasts. Further studies are warranted to assess the immune-modulating capability and anti-tumor activity of selective DDR1 inhibition in benign and malignant breast disorders.

## Materials and Methods

### Cell Culture and 3D collagen hydrogel formation

Immortalized human mammary fibroblasts, RMF-EG, were initially derived from primary human breast fibroblasts obtained from reduction mammoplasty tissues as described previously^31^. RMF-EG were cultured in DMEM (Corning 10017CV) with 10% FBS (Gibco) and 1% antibiotic/antimycotic (Corning MT30004CL). Collagen hydrogel 3D culture and contraction assays were performed as described previously^32^. Briefly, RMF-EG were seeded in rat tail collagen type 1 (Millipore Sigma 08115) hydrogels at varied cell number and collagen concentrations in 24 well ultra-low attachment surface plates (Corning 3473). Media without or without DDR1 inhibitor (DDR1-IN-1 [Tocris 5077] or 7rh [Tocris 5860] was added and gel contraction quantified at 8-16 hours. For collagen stimulation assays, RMF-EG were seeded into 6 well tissue culture plates (Corning 3516) with or without the addition of rat tail collagen type I to a final concentration of 0.05 mg/mL. For the TGFβ1 activation assays, RMF-EG 100,000 cells were seeded in 500uL collagen I gels (1mg/mL) or on tissue culture plate, exposed to recombinant human TGFβ1 (Thermo Fisher 7754BH005) 2ng/ml for 3 days, followed by treatment with DDR1i 1 μM for 24 hours.

### RNA extraction and RT-qPCR

Total RNA was isolated using TRIzol reagent (ThermoFisher 15596026) and RNeasy Mini Kit (Qiagen 74106). cDNA synthesis was performed using the iScript cDNA synthesis kit (Bio-Rad 1708891). qPCR assays were performed with the CFX Connect Real-Time PCR detection system (Bio-Rad) and iTaq Universal SYBR Green SuperMix (Bio-Rad 1725124). Each condition for the tested genes were repeated three times in triplicate. Gene expression analysis was performed using ddCq method with relative gene expression normalized to that of *GAPDH*. Mouse primer pairs used: *IL6*F-ACAAAGCCAGAGTCCTTCAGAG, R-GTGAGGAATGTCCACAAACTGA; *MCP*1: F-TTTTGTCACCAAGCTCAAGAGA; R-ATTAAGGCATCACAGTCCGAGT; *GAPDH* F-GATGACATCAAGAAGGTGGTG; R-GGTCCAGGGTTTCTTACTCCTT; *Mmp9* F-CAGCCGACTTTTGTGGTCTTC; R-CGGTACAAGTATGCCTCTGCCA; *Col1a1* F-CCCTGGTCCCTCTGGAAATG; R-GGACCTTTGCCCCCTTCTTT, *Col1a4* F-AAAGGCTCTCCGGGTTCAAT, R-CCGATGTCTCCACGACTAC; *Cxcl1* F-CCCAAACCGAAGTCATAGCCA, R-CTCCGTTACTTGGGGACACC; *Cxcl12* F-CCTTCAGATTGTTGCACGGC; R-CTTGCATCTCCCACGGATGT. Human primer pairs used: *SMA* F-CAGGGCTGTTTTCCCATCCAT, R-GCCATGTTCTATCGGGTACTTC; *FAP* F-AATGAGAGCACTCACACTGAAG, R-CCGATCAGGTGATAAGCCGTAAT; *IL6* F-AAGCCAGAGCTGTGCAGATGAGTA, R-TGTCCTGCAGCCACTGGTTC; *GAPDH* <u>F</u>-GAGTCAACGGATTTGGTCGT, R-TTGATTTTGGAGGGATCTCG; *IFNG* F-TCGGTAACTGACTTGAATGT, R-TCGCTTCCCTGTTTTAGCTC; *COL1A1* F-GACAGAGGCTCACAAGGTGAA, R-GTGGACCCATAGGACCGATG; *IL6* F-AAGCCAGAGCTGTGCAGATGA, R-TGTCCTGCAGCCACTGGTTC; *IL1B* F-CTCGCCAGTGAAATGATGGCT, R-GTCGGAGATTCGTAGCTGGAT; *IL8* F-TTTTGCCAAGGAGTGCTAAAGA, R-AACCCTCTGCACCCAGTTTTC; *MCP1* F-GAGAGGCTGAGACTAACCCAGA, R-ATCACAGCTTCTTTGGGACACT,

### Multiplex Cytokine Array

Conditioned media was generated by culturing RMF-EG in 3D hydrogels for 72 hours followed by the addition of serum free DMEM for 24 hours. Cell supernatants were centrifuged at 300g x 5 mins, filtered using a 0.22 µm membrane filter, and stored at −80C. Cytokine profiling of cell supernatants was performed using Human Cytokine Array C5 (RayBiotech AAH-CYT-5-2) and imaged using the ChemiDoc MP Imaging system (Bio-Rad). Relative protein expression was determined per manufacturer recommendations.

### In Vivo

Eight-week-old female C57BL/6J mice (Jackson Laboratories) were administered 7rh DDR1 inhibitor (Tocris 5860) 25 mg/kg or vehicle control (1% carboxymethylcellulose [Fisher Scientific C5678], 0.25% Tween 80 [Millipore Sigma P1754]) via oral gavage for 14 days. Mice were sacrificed followed by collection of inguinal/thoracic mammary glands and spleens for RT-qPCR, immunohistochemistry, and flow cytometry analysis. Inguinal lymph nodes were dissected away from inguinal mammary glands prior to RNA extraction and single cell suspension formation. RNA was extracted using a TissueLyser LT (Qiagen) and RNEasy MiniKit (Qiagen 74106), with RT-qPCR performed as described above. Inguinal mammary glands were fixed in 10% neutral buffered formalin (Millipore Sigma HT501128) followed by paraffin-embedding and picrosirius red staining as performed by the Tufts Animal Histology Core. Collagen deposition was assessed by quantifying the amount of picrosirius red staining in a minimum of 8 fields at 10x objective for each mammary gland section using ImageJ software, version 1.52a^33^.

### Flow cytometry

To form single cell suspensions for flow cytometry analysis, mammary glands were finely chopped and digested for 30-60 minutes at 37°C using a using a digestion mixture of collagenase 1.5 mg/mL and hyaluronidase 125 unit/mL in DMEM/F12 (Corning MT10090CV), 5% FBS (Gibco), insulin 10 µg/mL (Millipore Sigma I3536), mouse endothelial growth factor 5ng/mL (Millipore Sigma E5160), and hydrocortisone 0.5 µg/mL (Millipore Sigma), followed by the addition of RBC lysis buffer (HybriMax Sigma R7757), TrypLE Express Enzyme (Gibco 12604013), and DNase 0.1 mg/mL (Roche 1010415900). To form single cell suspensions from spleens, a mechanical digestion method was used. Spleens were homogenized using a 100µm strainer followed by the addition of RBC lysis buffer. The digestion mixturex were filtered through 100 µm and 40 µm filters and suspended in 1% BSA (Rockland BSA50) and 1 mM EDTA in phosphate buffered saline. Cells were incubated for 30 minutes at 4°C with myeloid or lymphoid antibody panels as previously described^28^; the following antibodies were used: CD45.2-APC-Cy7 (BioLegend 109823; 1:200), CD11b-PE (BioLegend 101207; 1:2400), CD11c-APC (BD 561119; 1:400), Ly6C-BV421 (BD 562727; 1:800), Ly6G-PE-Cy7 (BioLegend 127617; 1:2400), CD206-FITC (BioLegend 141703; 1:800), CD49b-FITC (BD 561067; 1:800), CD45.2-PE (BioLegend 109807; 1:400), TCRβ-PE-Cy7 (BioLegend 109221; 1:400), CD19-APC (BioLegend 115511; 1:800), CD8-BV711 (BioLegend 100747; 1:800), CD4-APC-Cy7 (BioLegend 100413; 1:400). Propidium iodide (1:5000) was added to each antibody panel for viability assessment. Flow cytometry was performed on a BD LSR-II flow cytometer at the Tufts Flow Cytometry Core. UltraComp eBeads Plus Compensation Beads (Invitrogen 01333341) were used for single-color compensation controls. Flow cytometry data were collected using FACS Diva, version 6.2. Flow cytometry compensation and data analysis was performed on FlowJo software v10.7.1. All analyses were performed in compliance with MiFlowCyt standards^34^.

## Acknowledgments

We gratefully acknowledge Tufts Comparative Medicine Services (CMS) for *in vivo* supportive services. We thank Nathan Li of Tufts Animal Histology Core for tissue processing and special stains. We thank Stephen Kwok and Allen Parmelee of the Tufts University Flow Cyometry Core for assistance with flow cytometry analysis. This research was supported by the Breast Cancer Research Foundation.

## Supporting Information

**Figure S1 -.**
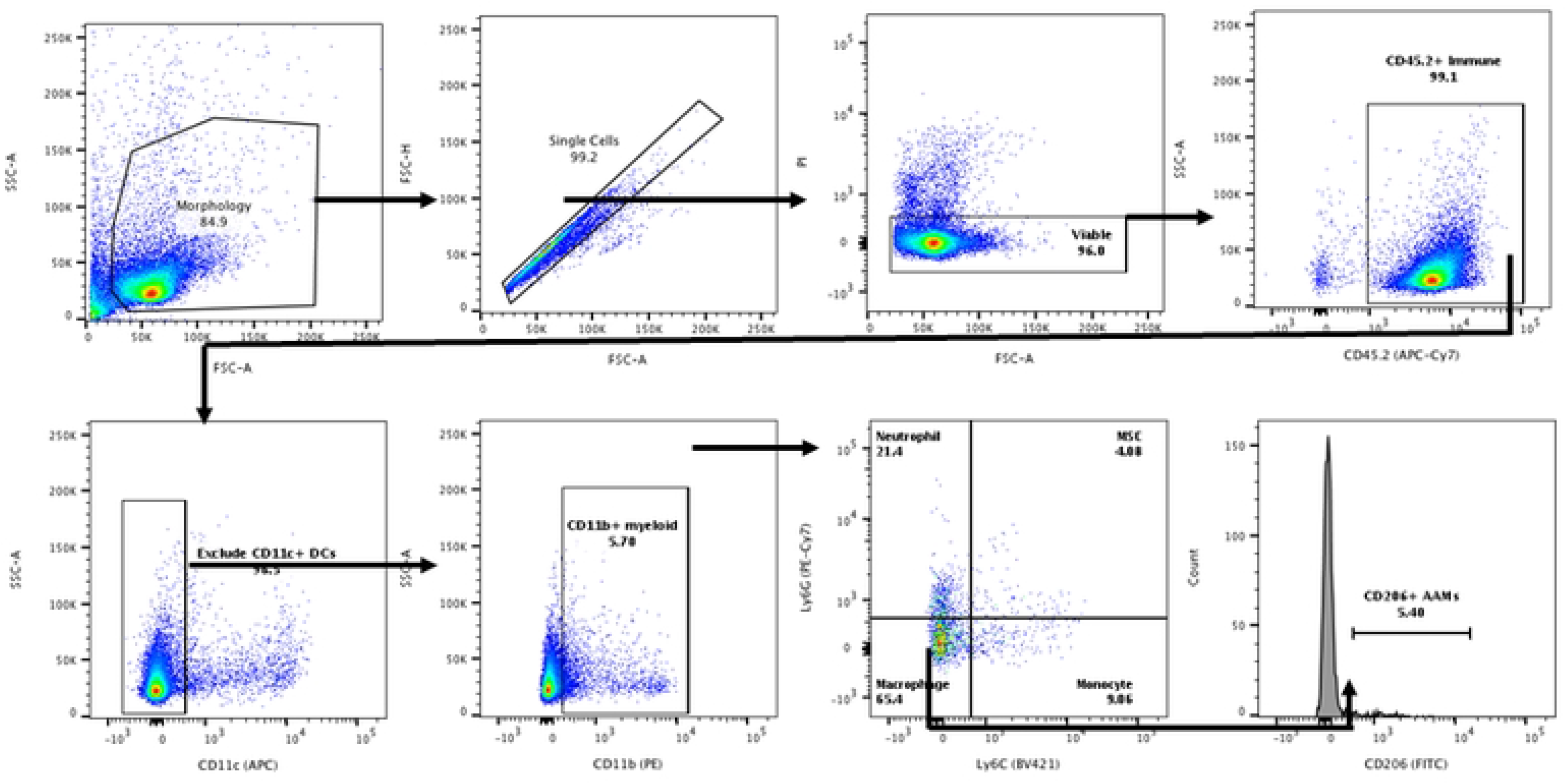
Gating method for flow cytometry assessment of myeloid immune cell populations.

**Figure S2 -.**
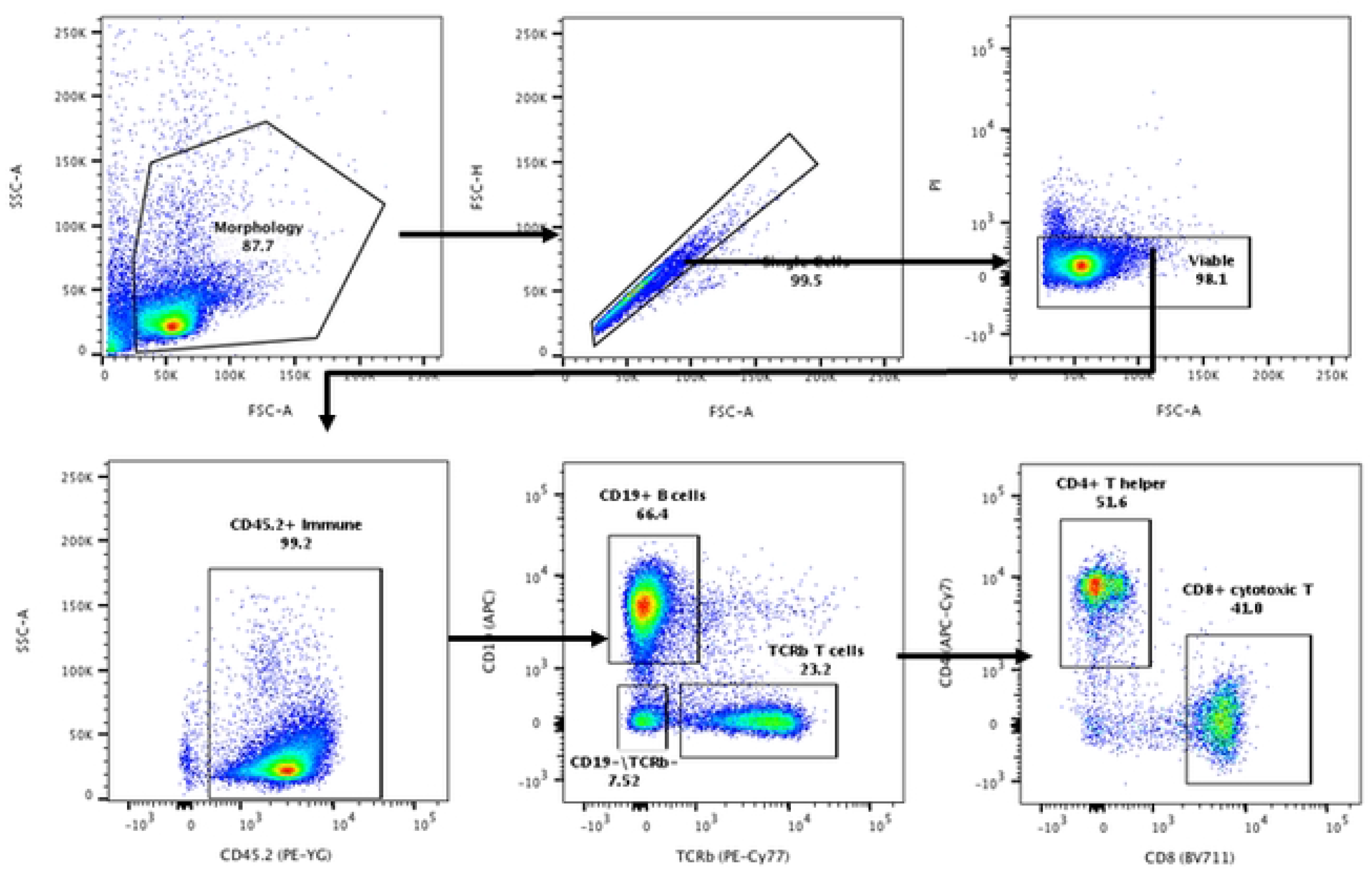
Gating method for flow cytometry assessment of lymphoid immune cell populations.

**Figure S3 -.**
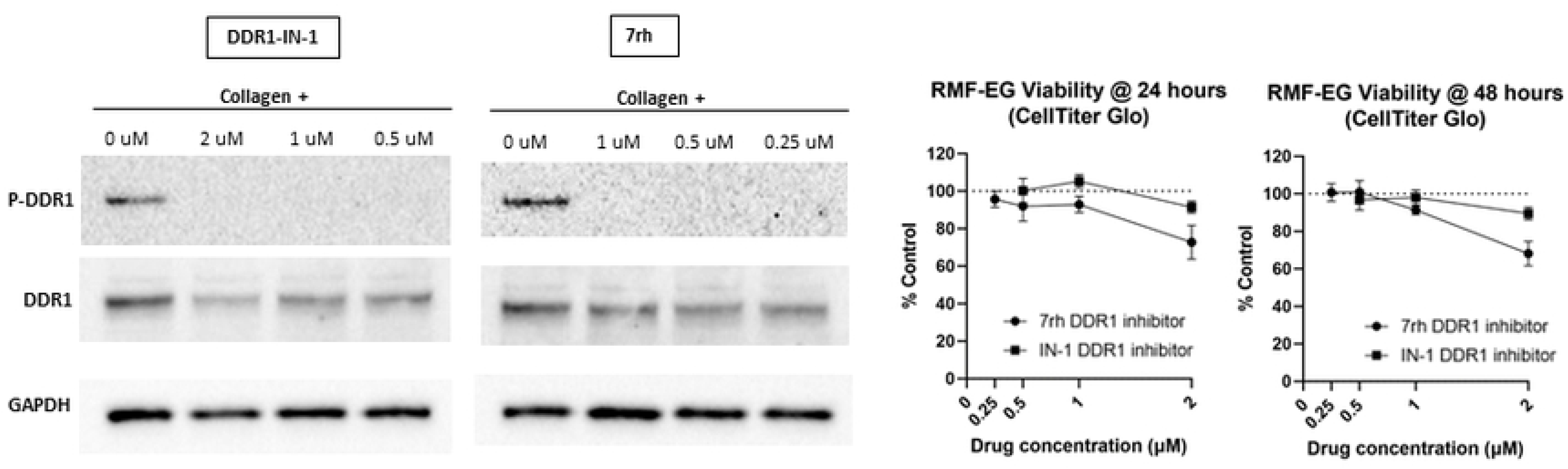
Dose titration of DOR1 inhibitors to assess cell viability and effective DDR1 inhibition.

**Figure S4 -.**
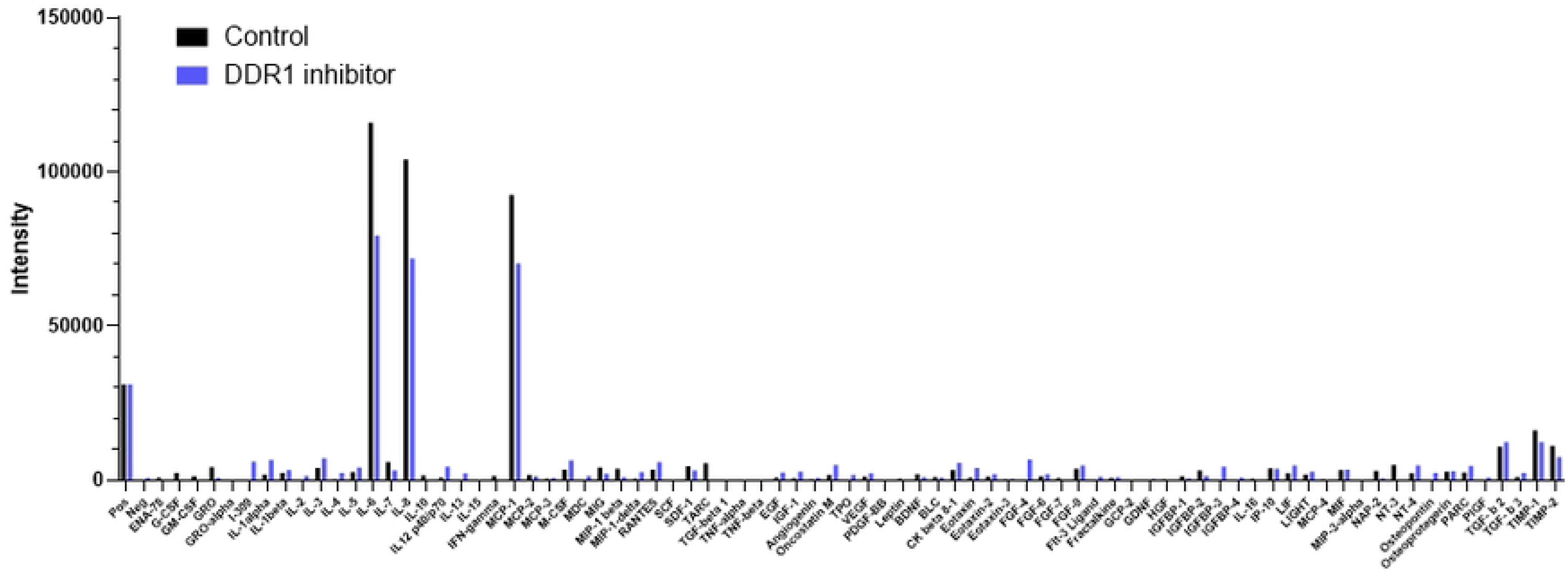
Human multiplex cytokine array densitometry analysis of RMF-EG cultured in 3D collagen hydrogel, total cytokine assessment.

**Figure S5 -.**
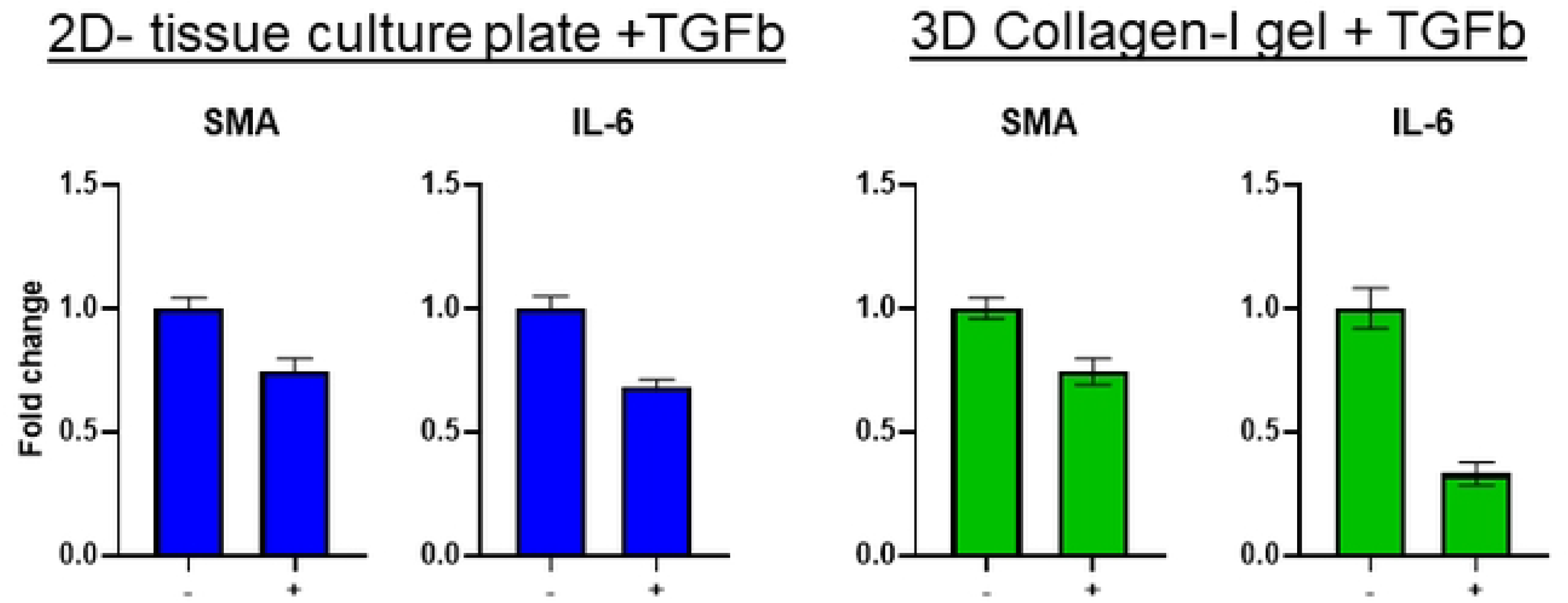
Assessment of SMA and IL-6 expression in RMF-EG cultured with recombinant TGFβ1 in the 2D or 3D context with (+) or without(−) OOR1i.

**Figure S6 -.**
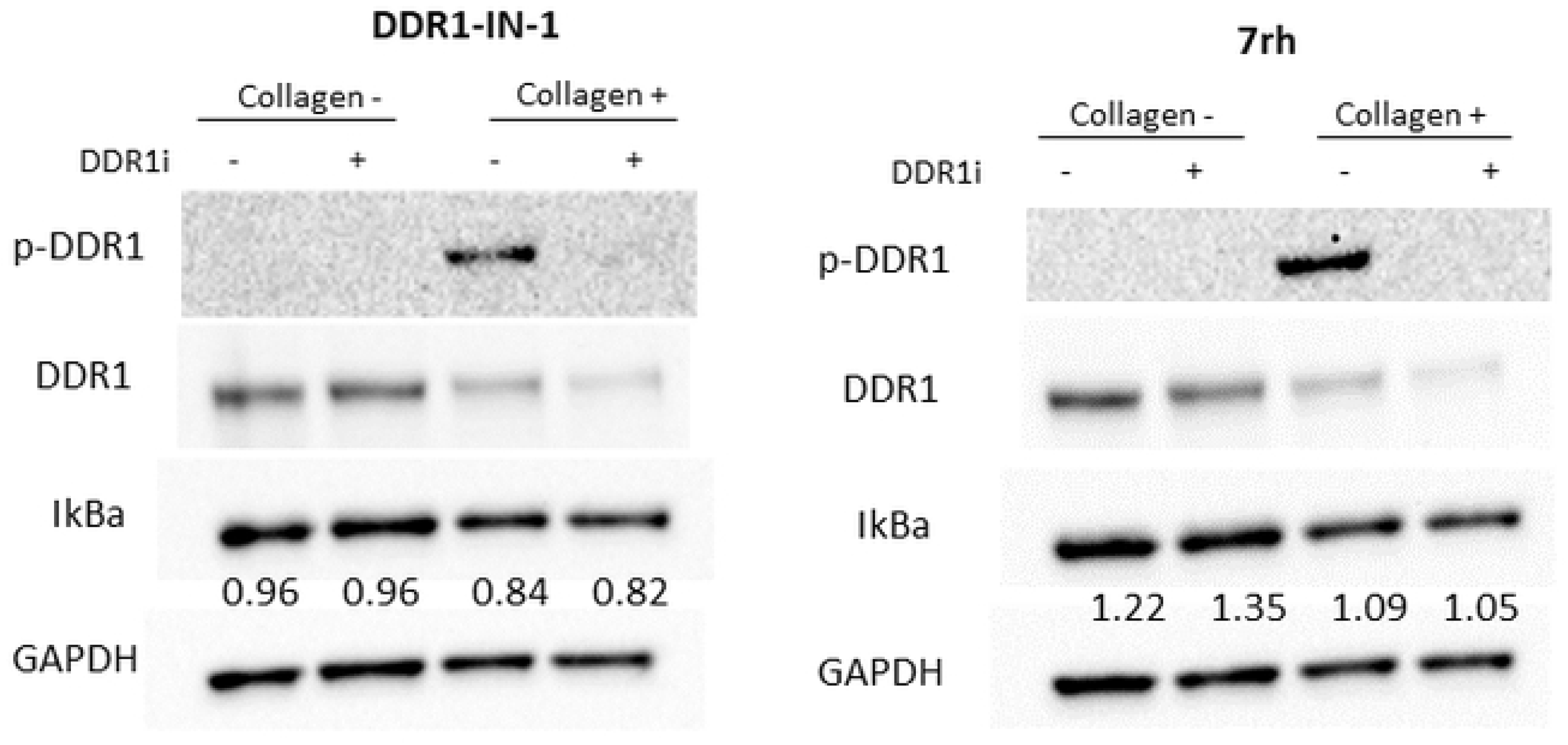
Effect of DDR1ion NF-κB signaling in RMF-EG.

